# Novel cholera toxin variant and ToxT regulon in environmental *Vibrio mimicus* strains: potential resources for the evolution of *Vibrio cholerae* hybrid strains

**DOI:** 10.1101/395061

**Authors:** Sucharit Basu Neogi, Nityananda Chowdhury, Sharda Prasad Awasthi, Masahiro Asakura, Zahid Hayat Mahmud, Mohammad Sirajul Islam, Atsushi Hinenoya, Gopinath Balakrish Nair, Shinji Yamasaki

## Abstract

Atypical El Tor strains of *Vibrio cholerae* O1 harboring variant *ctxB* genes of cholera toxin (CT) are gradually becoming a major cause of recent cholera epidemics. *Vibrio mimicus* occasionally contains virulence factors associated with cholera, e.g., CT, encoded by *ctxAB* on CTXФ genome; and TCP, the CTXФ-specific receptor. This study carried out extensive molecular characterization of CTXФ and ToxT regulon in *ctx*^*+ve*^ strains of *V. mimicus* isolated from the Bengal coast. Southern hybridization, PCR, and DNA sequencing of virulence related-genes revealed the presence of an El Tor type CTX prophage (CTX^ET^) carrying a novel *ctxAB*, tandem copies of environmental type pre-CTX prophage (pre-CTX^Env^), and RS1 elements, which were organized in an array of RS1-CTX^ET^-RS1-pre-CTX^Env^-pre-CTX^Env^. Additionally, a novel variant of *tcpA* and *toxT* respectively, showing clonal lineage to a phylogenetic clade of *V. cholerae* non-O1/O139, was identified. The *V. mimicus* strains lacked the RTX and TLC elements, and *Vibrio* seventh pandemic islands of the El Tor strains, but contained five heptamer (TTTTGAT) repeats in *ctxAB* promoter region like some classical strains of *V. cholerae* O1. PFGE analysis showed all the *ctx*^+ve^ *V. mimicus* strains were clonally related. However, their *in vitro* CT production and *in vivo* toxigenecity were variable, which could be explained by differential transcription of virulence genes along with ToxR regulon. Taken together, our findings strongly suggest that environmental *V. mimicus* strains act as potential reservoir of atypical virulence factors, including variant CT and ToxT regulon, and may contribute to the evolution of *V. cholerae* hybrid strains.

**IMPORTANCE:** Natural diversification of CTXФ and *ctxAB* genes certainly influences disease severity and shifting patterns in major etiological agents of cholera, e.g., the overwhelming emergence of hybrid El Tor variants, replacing the prototype El Tor strains of *V. cholerae*. This study showing the occurrence of CTX^ET^ comprising a novel variant of *ctxAB* in *V. mimicus* points out a previously unnoticed evolutionary event, independent to that of the El Tor strains of *V. cholerae*. Identification and cluster analysis of the newly-discovered alleles of *tcpA* and *toxT* indicates their horizontal transfer from an uncommon clone of *V. cholerae*. The genomic content of ToxT regulon, and tandemly arranged multiple pre-CTXФ^Env^ and a CTXФ^ET^ in *V. mimicus* probably act as salient raw materials inducing natural recombination among the hallmark virulence genes of hybrid *V. cholerae* strains. This study will facilitate deeper understanding of the evolution of new variant CT and ToxT regulon, influencing cholera epidemiology.

## INTRODUCTION

**V***ibrio mimicus* is genetically and ecologically very similar to *Vibrio cholerae*, the cholera bacterium and share similar environmental niche in freshwater and estuarine ecosystems, particularly in the tropical region like the Bengal delta. *V. mimicus* is known to be associated with sporadic cholera-like diarrhea cases. Despite a lot of efforts in hygiene promotion and therapeutic advances, cholera continues to pose as a major health problem worldwide, accounting for millions of episodes and thousands of deaths, with ca. 132,000 cases in 2016 reported to the World Health Organization (http://www.who.int/gho/epidemic_diseases/cholera/en/). The principal pathogenic factor instigating the disease is the cholera toxin (CT), encoded by the *ctxAB* operon, predominantly found in *V. cholerae* strains belonging to the O1 and O139 serogroups, and occasionally a few non-O1/non-O139 serogroups. Among the seven known cholera pandemics, the current seventh pandemic since 1961 is caused by the El Tor biotype of *V. cholerae* O1 while its classical biotype was associated with the sixth pandemic. In Bangladesh, the classical cholera re-emerged in 1983, later receded by the rise in El Tor cholera, and is believed to be extinct since 1993. However, since the last decade, hybrid El Tor strains producing classical-CT are the dominant cause of epidemic and endemic cholera replacing the prototype El Tor strains that produce El Tor CT (1). Occurrences of such type of variant El Tor strains have also reported to spread in many countries in Asia, Africa, and in Haiti (2, 3, 4). This indicates a cryptic existence of the variant or classical *ctxB*, and variant CTXФ in environmental reservoirs, yet mostly unexplored. *In vitro* experiments have shown that CTXФ can infect certain *V. mimicus* strains (5). In line with this, occurrence of *ctxAB* among *V. mimicus* strains, although isolated rarely, in Bangladesh, India, Japan and the United States, attests the hypothesis of inter-species genetic exchange (6, 7, 8, 9).

The *ctx*AB operon encoding the A and B subunits of CT is a part of the genome of CTXФ, a filamentous bacteriophage. The precursor form of the CTXФ, pre-CTXФ, does not carry the *ctxAB* genes (8). Before this study, a total of 13 genotypes of *ctxB* have been distinguished based on single nucleotide polymorphisms (SNPs) at 10 loci of this toxigenic factor (Table 2). Notably, the *ctxB* genotypes 1 and 2 are typical for all classical strains and El Tor strains from Australia, respectively, while genotypes 3 and 7 are featured among the pandemic El Tor, and the Haitian variant strains. *V. cholerae* O1 El Tor strains are also characterized by the presence of TLC (Toxin linked cryptic) element and repeat in toxin (RTX) genes in the flanking region of CTX prophage, and two large genomic islands, termed as *Vibrio* Seventh Pandemic Islands (VSP-I and VSP-II) (10). Other known virulence factors of *V. cholerae*, particularly of the non-O1/non-O139 strains, include heat-stable enterotoxin (encoded by *stn*), type III secretion system (*vcsN2*), and cytotoxic cholix toxin (*chxA*) (11, 12). Natural recombination events, compounded with the integration of phages contribute to evolution of genes, especially those related to virulence and ecological fitness (13).While persisting in the aquatic environment *V. cholerae* and *V. mimicus* interact with diverse phages, and a portion of their populations, harboring selective receptor, can integrate toxigenic phages into their genome.

**Table 1.**
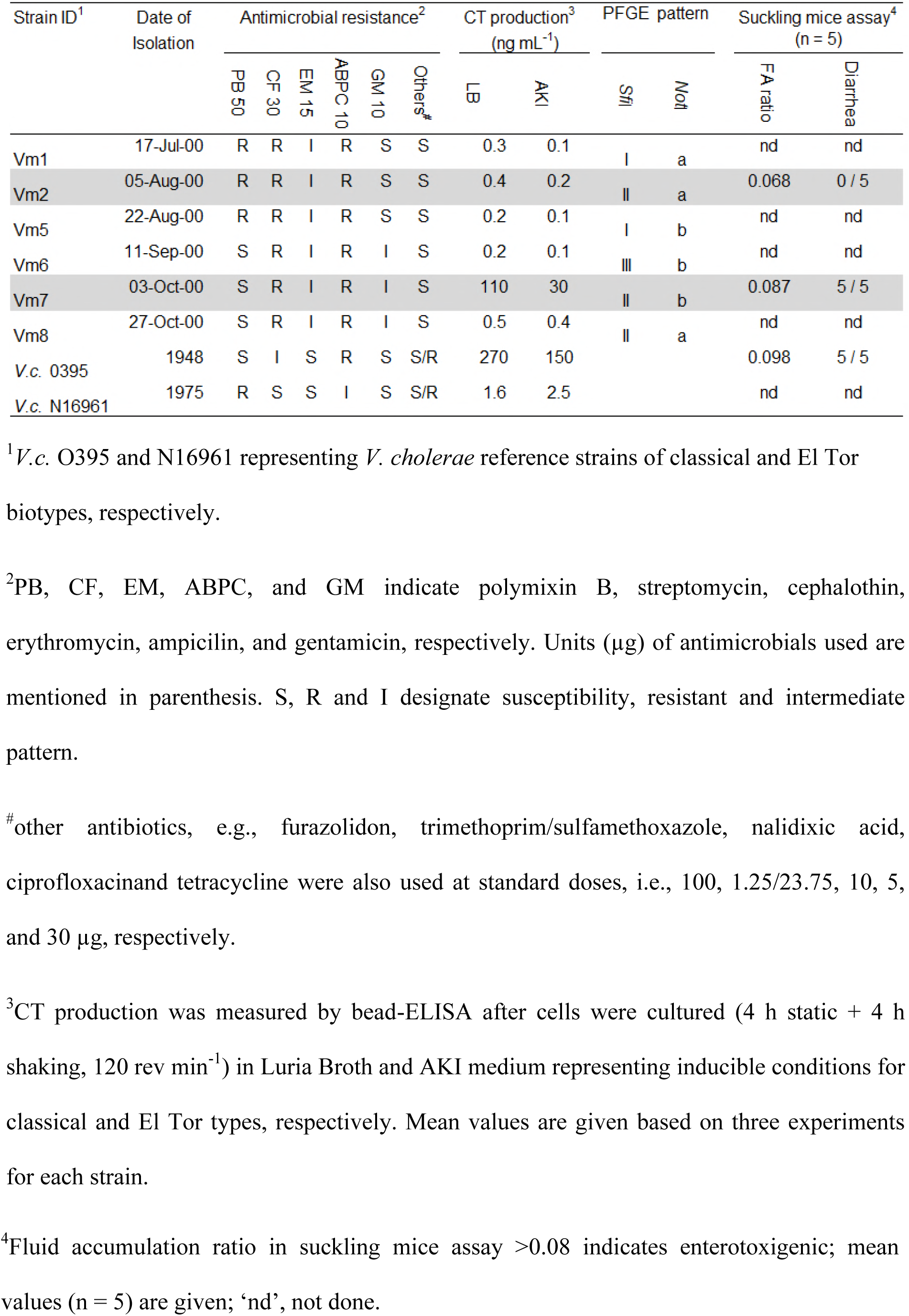
Antimicrobial susceptibility, cholera toxin production, PFGE pattern and enterotoxigenicity in *ctx*^+ve^ *V. mimicus* strains

**Table 2.** Comparative diversity in *ctxAB* gene among *V. mimicus* and *V. cholerae*

| Strains<sup>1</sup> | Isolation |  | *ctxA* (aa positions) |  |  |  |  | *ctxB* (aa positions)<sup>2</sup> |  |  |  |  |  |  |  |  |  | Designated Genotype | Reference |
| --- | --- | --- | --- | --- | --- | --- | --- | --- | --- | --- | --- | --- | --- | --- | --- | --- | --- | --- | --- |
|  | Country | Year | 46 | 190 | 198 | 226 | 255 | 20 | 24 | 28 | 34 | 36 | **39** | 46 | 55 | 67 | **68** |  |  |
| VC O1, CL, O395 (CP000627) | India | 1948 | S | R | I | V | K | H | Q | D | H | T | H | F | K | A | T | 1 | [1] |
| VC O1, Australia ET | Australia |  | - | - | - | - | - | H | Q | D | H | T | H | L | K | A | T | 2 | [39] |
| VC O1, ET, N16961 (NC_002505) | Bangladesh | 1975 | S | R | I | V | K | H | Q | D | H | T | Y | F | K | A | I | 3 | [39] |
| VC O139 (FJ821557) | Bangladesh | 1998 | - | - | - | - | - | H | Q | D | H | T | Y | F | K | A | T | 4 | [39] |
| VC O139 (FJ821556) | Bangladesh | 2005 | - | - | - | - | - | H | Q | A | H | T | H | F | K | A | T | 5 | [39] |
| VC O139 (FJ821581) | Bangladesh | 2007 | - | - | - | - | - | H | Q | D | P | T | Y | F | K | A | T | 6 | [39] |
| VC O1 (EU496273, L19089) | India, Haiti | 2007, 2010 | - | - | - | - | - | N | Q | D | H | T | H | F | K | A | T | 7 | [39] |
| VC O27 (AF390572) | Japan | 1996 | S | R | I | V | E | H | H | A | H | T | H | F | K | A | T | 8 | [39] |
| VC O37 (D30052) | Sudan | 1968 | N | R | I | V | K | H | Q | D | H | T | H | L | N | A | T | 9 | [39] |
| VC O1 (EU932878) | Zambia | 1996 |  |  |  |  |  | H | Q | D | P | T | Y | F | K | A | I | 10 | [54] |
| VC O1 (EU932881) | Zambia | 2003 |  |  |  |  |  | H | Q | D | P | T | H | F | K | A | T | 11 | [54] |
| VM (ACYV01000039) | USA | 1990 | N | I | I | I | K | H | Q | D | H | T | H | L | K | A | T | 12 | [19] |
| VC O1 (SH65928) | China | 1965 | - | - | - | - | - | H | Q | D | H | A | Y | L | N | A | T | 13 | [55] |
| **VM (This study)** | Bangladesh | 2000 | **N** | **I** | **V\*** | V | K | H | Q | D | H | T | H | L | K | **E\*** | T | **14** | This study |

<sup>1</sup>VC, VM, CL and ET represent *V. cholerae*, *V. mimicus*, classical and El Tor, respectively. Known serogroups of *V. cholerae* strains are shown, accession number of the gene sequences are given in parenthesis.
<sup>2</sup>The deduced amino acid (aa) positions are indicated by vertical numbering; bolded 39 and 68 positions bear the amino acid markers, differentiating classical and El Tor type *ctxB* gene.
\*Unique change in deduced amino acid of *ctxAB* in *V. mimicus* strains of this study.

The CTXФ genome (∼ 6.9 kb) contains core and RS2 regions. The core region includes genes involved in phage morphogenesis and CT production, including *ctxAB, zot*, and *orfU*. The RS2 region contains genes required for replication (*rstA*), integration (*rstB*) and regulation (*rstR*) of CTXФ (14). Moreover, the upstream promoter of *ctxAB* possesses heptamer repeats, considered as evolutionary signature, while its downstream intergenic region contains site for CTXФ integration, mediated by XerC and XerD recombinases (15). In El Tor strains, the prophage DNA is flanked by a genetic element known as RS1, which is a satellite phage (16). In comparison to RS2, the RS1 additionally contains *rstC* that ecodes an anti-repressor of *rstR* and promotes transmission of RS1 and CTXФ (17). In *V. cholerae* strains, presence of both CTX prophage and RS1 element, as solitary and multiple copies with diverse arrays of genetic organization, have been documented (18). Based on nucleotide sequence polymorphism in its several genes, including *rstR* and *orfU* (gIII^CTX^), the CTX prophage can be differentiated into several types such as classical, El Tor, Calcutta and environmental (19). Among the El Tor variant or hybrid strains, two types of CTX prophages, one harboring classical *rstR* and classical *ctxB* (20) and the other containing El Tor *rstR* and classical *ctxB* (21) have been reported. Although extensive investigations have revealed nucleotide sequence polymorphism and diversity in the array of CTX prophages on *V. cholerae* genome (21) little is known for those of *V. mimicus* strains.

The transmission of CTXФ into a *Vibrio* strain relies on the presence of a specific cell surface type IV pilus receptor, termed as toxin co-regulated pilus (TCP), which also plays a vital role aiding colonization of *V. cholerae* in human or animal intestine (22). The TCP is located on the *Vibrio* Pathogenicity Island (VPI), and produced by the action of a cluster of genes, termed as TCP island. The major structural subunit of TCP is encoded by *tcpA*. The expression of CT and TCP is activated by ToxT, present on the TCP island, and is under the control of the ToxR regulon, comprising *toxR, toxS, tcpP*, and *tcpH* (23). Based on the nucleotide sequence polymorphism in *tcpA*, the TCP can be differentiated into several types, e.g., El Tor, classical, Nandi, and Novais (24). CTX/pre-CTX prophages and genes of VPIs are found scattered throughout environmental isolates of *V. cholerae* (25). Despite the absence of the classical biotype strains along with the classical CTX phage particle (1), the increasing occurrence of hybrid El Tor strains of *V. cholerae* O1 harboring variant *ctxB* genes is intriguing and requires detail exploration for their environmental reservoirs. Being genetically the closest species of *V. cholerae*, there is high possibility for the environmental *V. mimicus* strains to act as potential reservoir of virulence genes associated with cholera and diarrhea epidemics. However, our knowledge on the occurrence of genetic determinants of virulence, particularly cholera-like diarrhea, in environmental *V. mimicus* and their similarity to those of epidemic strains of *V. cholerae* is very limited.

In this study, several *ctx*^+ve^ *V. mimicus* strains isolated from estuarine surface waters in Bangladesh were analyzed to ascertain whether they can act as reservoirs of the CTXФ carrying *ctxAB* variant present in *V. cholerae* strains associated with recent epidemics. The objectives were to investigate (i) the molecular diversity of genetic elements within CTX prophage and TCP islands, (ii) *in vitro* CT production, (iii) *in vivo* fluid accumulation using suckling mouse model, and (iv) differential expression of ToxT regulon in these environmental *V. mimicus* strains. Comparison of these phenotypic and genetic traits to those of toxigenic *V. cholerae* would aid in better understanding the evolution of new variant CT and ToxT regulon.

## RESULTS

### Antimicrobial susceptibility

Among the 11 antimicrobials tested, all *ctx*^+ve^ *V. mimicus* strains examined in this study showed full resistance to ampicilin (10 μg) and cephalothin (30 μg), and intermediate resistance to erythromycin (15 μg). However, two types of antimicrobial resistance pattern were observed based on the resistance to polymixin B (50 μg) and gentamicin (10 μg) (Table 1). Three out of six *V. mimicus* strains showed full resistance to polymixin B (50 μg), while the other three strains showed intermediate resistance to gentamicin (10 μg).

### PFGE based screening for genomic relatedness

PFGE of the undigested gDNA showed that the *ctx*^+ve^ environmental *V. mimicus* strains possessed ca. 2.9 and 1.3 Mbp of large and small chromosomes, respectively, which were similar to *V. mimicus* type strain (ATCC33539^T^) but different from the classical (O395), El Tor (N16961) and non-O1/non-O139 (VCE233) strains of *V. cholerae*. PFGE analysis of *Not*I- and *Sfi*I-digested gDNA showed 0 to 3 band differences among the *ctx*^+ve^ *V. mimicus* strains. According to 26 Tenover *et al*. (1995), these strains were clonal in origin. However, comparison of the PFGE bands with two enzymes could reveal a total of five subtypes (Table 1). In case of *Sfi*I, three patterns (designated as I, II and III, Fig. S1) could be assigned, but only two patterns (designated as a and b, Fig. S1) were observed in case of *Not*I. Taken together, four subtypes (patterns Ia, Ib, IIb and IIIb) were present in four out of six strains. The remaining two strains did not show any difference (pattern IIa) even after digestion with both the enzymes.

### Occurrence of the major virulence factors

Among the hallmark genes associated with the toxigenic *V. cholerae* strains, several of them related to CT production were detected in the *V. mimicus* strains used in this study. Colony blot hybridization using ^32^P-labelled probes for virulence related-genes showed the presence of *ctxA* and *zot* of CTXФ, *rstC* of RS1 element, and *tcpA* of VPI. However, these strains did not harbor genes representing VSP I and II. They were also negative for TLC and RTX elements, which are commonly present in the flanking region of CTXФ in *V. cholerae* El Tor and their hybrid strains. No other major toxigenic factors of *V. cholerae*, namely, *vcsN2, chxA*, and *stn* were detected in the *ctx*^+ve^ *V. mimicus* strains.

### Characteristics of *ctxAB* and CTXФ associated genetic elements

Based on the results of MAMA-PCR, all the *V. mimicus* strains contained classical type of *ctxB.* Sequence analysis of the entire *ctxB* observed that the gene was identical in all six *V. mimicus* strains. Although showing signature changes at the 39th (tyrosine to histidine) and 68th (isoleucine to threonine) positions, similar to classical *ctxB* genotype 1, the *V. mimicus ctxB* had additional non-synonymous substitutions conferring subtle changes in the deduced amino acids at positions 46 (phenylalanine to leucine) and 67 (alanine to glutamic acid) (Table 2). Comparative analysis with other known *ctxAB* genotypes reported among *V. mimicus* and *V. cholerae* revealed the existence of a new *ctxB*, designated as genotype 14. This genotype was almost similar to genotype 12, reported from a *V. mimicus* strain, but differed at amino acid position 67. In all other genotypes of *ctxB* the 67^th^ position encoded alanine, but all the *ctxB* sequences of *V. mimicus* strains in this study detected glutamic acid at this position, thus unique for this novel genotype 14.

Sequencing analysis of *ctxA* of the environmental *V. mimicus* strains also observed alterations from the canonical gene and all available sequences in GenBank. This novel *ctxA* differed from that of the reference El Tor and classical strains with three amino acids at positions 46, 190 and 198, while the highest similarity was observed with *ctxA* of a *V. mimicus*, characterized with *ctxB* genotype 12 (Table 2). A unique change at amino acid position 198, with alteration of isoleucine to valine, of *ctxA* in the examined *V. mimicus* strains was noteworthy.

The presence of RS1 element was confirmed by the *rstC* gene-based PCR (Fig. S2), followed by sequencing analysis. The *rstC* gene in the environmental *V. mimicus* strains was identical to that of the reference El Tor strain. PCR-based genotyping of *rstR* showed the presence of two different alleles, one for El Tor (*rstR*^ET^) and the other for environmental (*rstR*^Env^), indicating the occurrence of multiple prophages, i.e., CTXФ^ET^ and CTXФ^ET^.

### Genetic organization of CTXФ associated elements

Southern hybridization of chromosomal DNA digested with *Bgl*I and *Bgl*II, which have single cutting site in CTXФ at *rstR*^ET^/ *rstR*^Env^ and *zot*, respectively, showed identical RFLP patterns for all *V. mimicus* strains. Size-wise comparative analysis of the bands detected by hybridization, using different probes, of the enzyme digested gDNA revealed similar results, i.e., presence of two copies of pre-CTXФ^Env^ prophages and a single CTXФ^ET^ prophage, in all six strains of *ctx*^+ve^ *V. mimicus*. Probing with *ctxA* and *rstR*^Env^ of the *Bgl*I-digested gDNA generated a single positive band (ca. 10.5 Kb), and two positive bands (ca. 6.5 and 10.5 Kb), respectively. These results indicated the presence of one copy of CTXΦ containing *ctxAB*, followed by a pre-CTXФ^Env^ (lacking *ctxAB*) with an adjacent RS1, and pre-CTXФ^Env^ without RS1 (Fig. 1, Fig. S3). Probing with *rstR*^Env^ and *rstR*^ET^ of the *Bgl*II-digested gDNA resulted one (ca. 18.8 kb), and three (ca. 3.5, 8.0 and 18.8 Kb) positive bands, justifying the adjacent locations of one RS1 followed by two pre-CTXФ^Env^ prophages, along with the preceding occurrence of another RS1 and El Tor type full length CTXΦ containing *ctxAB*. Hybridization with *rstC* probe justified the presence of two copies of RS1 element, one before the CTXФ^ET^ and the other preceding the adjacent CTXФ^Env^. However, there was no RS1 element between the two adjacent CTXФ^Env^ prophages. Altogether, an array of RS1-CTXФ^ET^-RS1-pre-CTXФ^Env^-pre-CTXФ^Env^ was deduced from the Southern hybridization analysis.

**Fig. 1.**
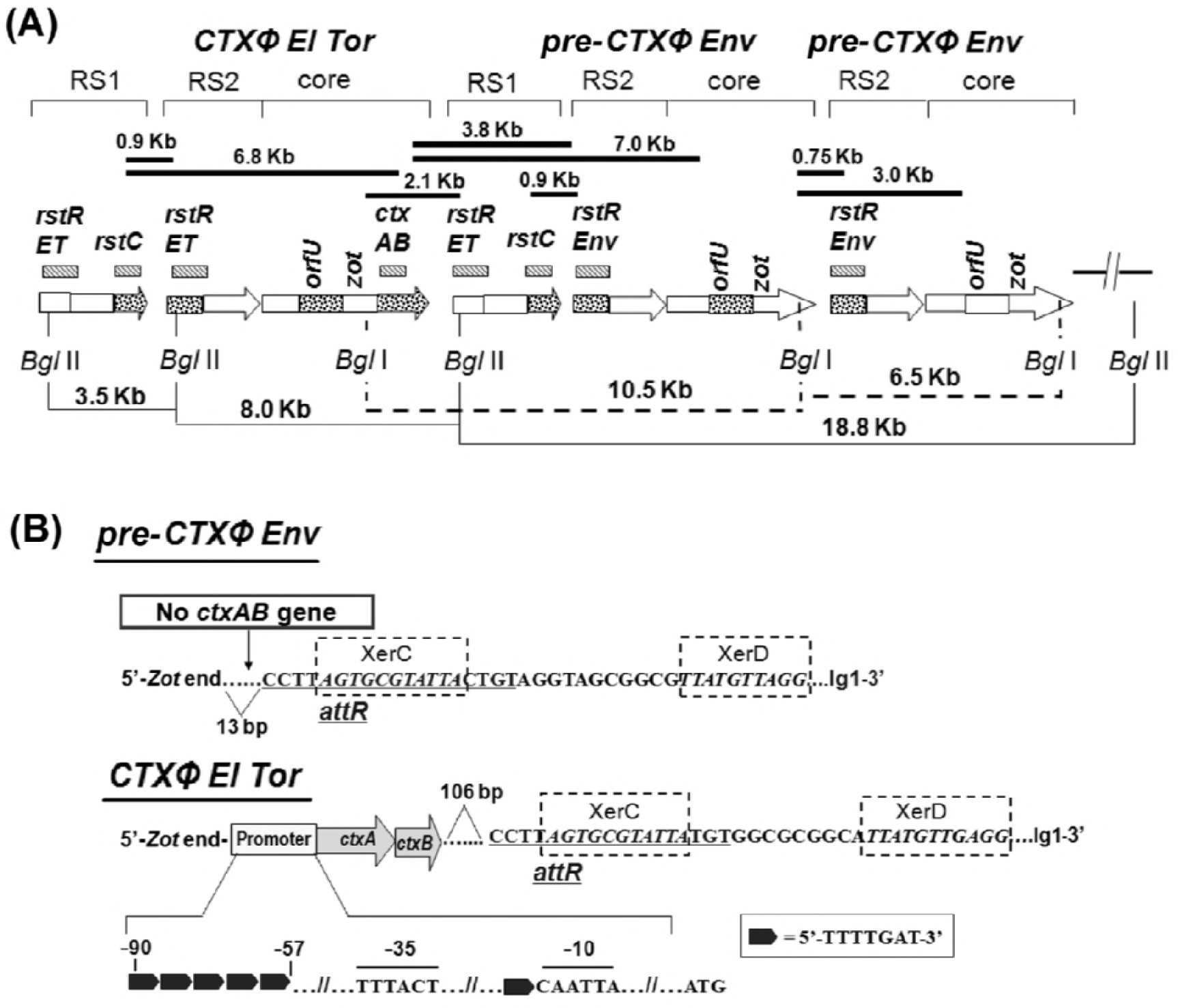
Organization of CTXФ^El^ ^Tor^, RS1 and pre-CTXФ^Env^ in *V. mimicus* strains. (A) Filled bars indicate PCR arrays used to check probable locations of genes and sizes of PCR products are given on top. Hashed bars indicate the genetic regions (names mentioned on top) used as probes for Southern hybridization after restriction digestion with *Bgl*I or *Bgl*II enzymes; arrows indicate RS1, RS2 or core prophage where dotted regions were analyzed by sequencing. Lines (filled and dotted) in the bottom show the distances between specific genetic locations determined by Southern hybridization analysis of the *Bgl*I- or *Bgl*II-digested genomic DNA using specific probes. (B) Region between *zot* and *rstR* in pre-CTXФ^Env^ and CTXФ^El^ ^Tor^ in *V. mimicus.* The *ctxAB* promoter of CTXФ^El^ ^Tor^ contains 5 heptamer (TTTTGAT) repeats, shown by filled black arrows, which is characteristic of classical type *ctxAB*. In the reference El Tor strain, N16961, the attR sequence is also located 106 bp downstream of *ctxAB*, followed by XerC and XerD.

To verify the hybridization results, PCR arrays using the allele-specific forward and reverse primers, with multiple combinations, of *rstC, rstR, ctxAB* and *orfU* genes (Fig. 1) were conducted. All the six *ctx*^+ve^ *V. mimicus* strains yielded similar amplicons after PCR using primers specific for different regions of CTX element. Size-wise comparison of the PCR amplicons observed concordance with the published genetic organization of El Tor strains of *V. cholerae*. The results of PCR walking were in concordance with hybridization results, confirming the presence of the RS1, *rstR*^ET^ in RS2 of the CTXΦ carrying *ctxAB*, and *rstR*^Env^ allele in RS2 of pre-CTXΦ element(s), which lacked the *ctxAB* operon (Fig. S2, Fig. S3). The flanking sequences of CTXФ^ET^ harbored both the RS1 and RS2 elements, similar to the reference *V. cholerae* O1 El Tor strain N16961. Sequencing analysis of several core and intergenic regions of CTX element (Fig.1), confirmed the tandem presence of three copies of the CTX element, including one intact CTXФ^ET^ and two pre-CTXФ^Env^ prophages lacking *ctxAB*.

### Genomic signatures in *ctxAB* promoter and intergenic sequences

The sequential integration of pre-CTXФ^Env^ (lacking *ctxAB*) and CTXФ^ET^ in *V. mimicus* strains isolated from the estuarine environment prompted sequencing analysis of intergenic regions, particularly *ctxAB* promoter and the prophage flanking regions to reveal genomic signatures associated with their lysogenic transformation. Forward and reverse primers of the adjacent genes, i.e., *zot, ctxA, ctxB* and *rstR*^ET^ for El Tor CTX prophage, and *zot* and *rstR*^Env^ for environmental pre-CTX prophage amplified the desired parts of *ctxAB* promoter and intergenic regions. Sequencing analysis of the El Tor CTX prophage showed that the promoter at the 5′- upstream of *ctxAB* contained 5 heptamer (TTTTGAT) repeat spanning between −90 and −57 bp, which is a characteristic of the classical type *ctxAB* promoter, while the RNA polymerase binding sites at −35 bp (TTTACT) and −10 bp (CAATTA) were conserved (Fig. 1). The 3′- end of *ctxAB* was characterized by attR sequences coupled with the XerC and XerD binding sites for CTXФ integration, which started 106 bp downstream similar to the reference El Tor strain N16961 of *V. cholerae*. On the other hand, the 3′-end of pre-CTX^Env^ prophage was characterized by a 13 bp gap between *zot* (the last gene of phage core region) and *attR* sequences, followed by XerC and XerD binding sites (Fig. 1).

### Sequence diversity in *orfU* of CTX prophage core region

Because the *orfU* gene is instrumental in studying the diversity in the core region of CTX prophage, this gene was PCR-amplified from *V. mimicus* strains and subjected to sequencing, followed by phylogenetic analysis. Interestingly, identical sequence homology among all the study strains and also within CTXФ^ET^ and pre-CTXФ^Env^ prophages was observed. Comparative analysis with the reference El Tor and classical strains of *V. cholerae* and a recently studied *V. mimicus*, of the deduced amino acid sequences of *orfU*, indicated that the environmental *V. mimicus* strains of this study did not completely match any of them rather they possessed nine unique changes with a total of 31 polymorphic sites observed within these strains. The highest similarity was observed with the reference El Tor strain N16961 of *V. cholerae* O1, which differed by 11 amino acids in *OrfU*. In comparison to OrfU of *V. mimicus* strains of this study, both the reference classical O395 strain of *V. cholerae* O1 and another *V. mimicus* strain differed, although not identical, by 27 amino acid substitutions. Thus, sequencing analysis indicated the presence of a variant OrfU (Fig. S4) in the genome of environmental *V. mimicus* strains. Phylogenetic analysis using partially available nucleotide sequences (702 of 1083 bp) of *orfU* in GenBank database showed this gene in the study strains did not cluster with the classical *V. cholerae* strains like the previously reported strains of *V. mimicus* (7). However, the *orfU* variant in environmental *V. mimicus* strains grouped into a cluster comprising of the El Tor, El Tor variant O1, O139, and several non-O1/non-O139 strains of *V. cholerae* (Fig. 2).

**Fig. 2.**
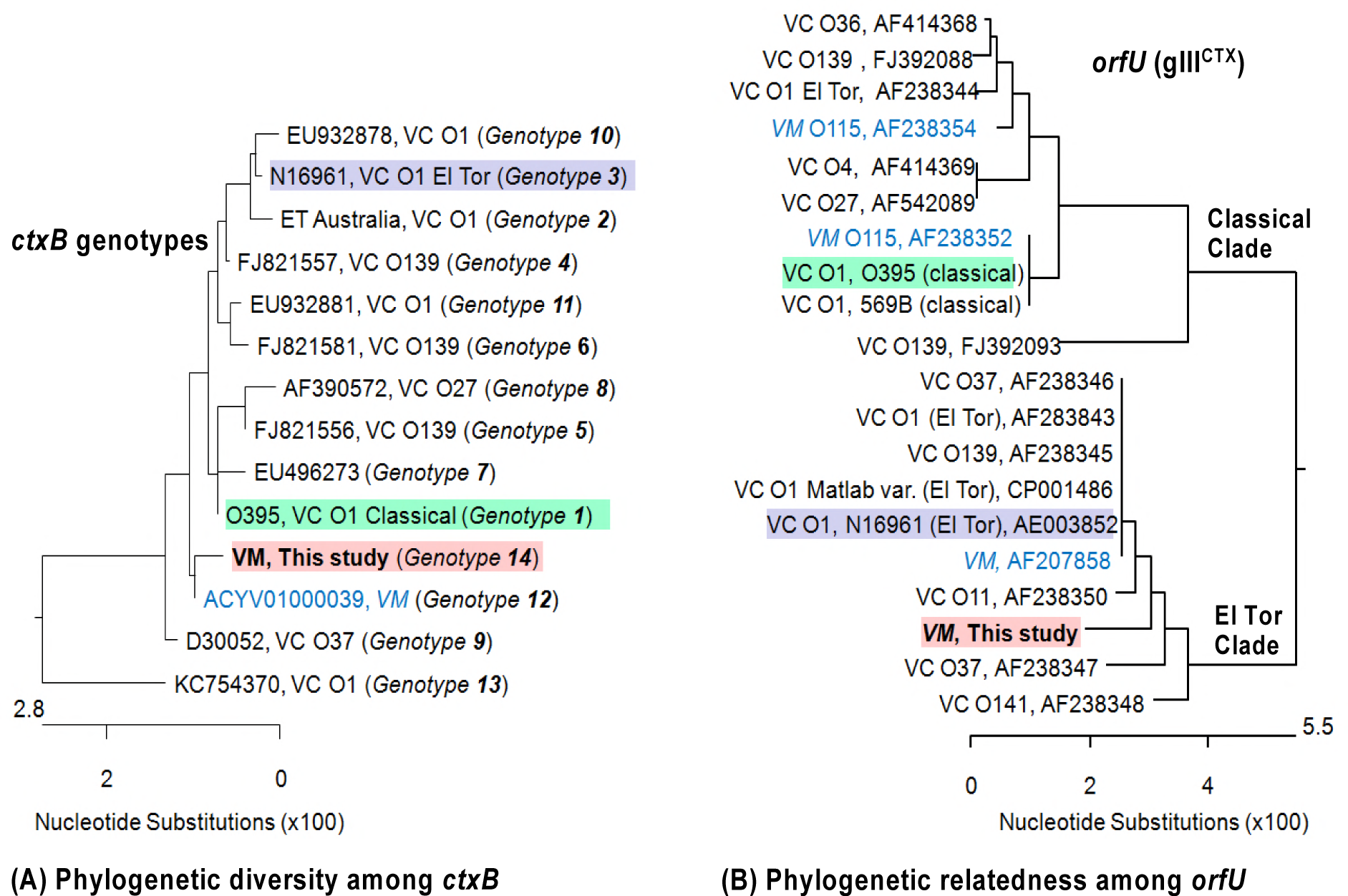
Genetic relatedness among *ctxB* and *orfU* genes of *V. mimicus* and *V. cholerae* strains. (A) The novel *ctxB* genotype 14 of *V. mimicus* showed closeness with both the genotypes 1 and 7, representing classical and Haitian strains, respectively. (B) The *orfU* of *V. mimicus* of this study showed close affiliation to the strains grouped into the El Tor clade.

### PCR, sequencing, and phylogenetic analysis of *tcpA* and *toxT*

PCR using a forward primer from the beginning part of the 5’-terminal conserved region (*tcpA*-*F*) and a reverse primer for El Tor or classical type of *tcpA* did not yield any amplicon. However, using *tcpA*-F and a reverse primer from *tcpQ* (*tcpQ*-R) yielded a 2.1-kb product for all the *ctx*^+ve^ *V. mimicus* strains. DNA sequencing showed that all of the strains had identical *tcpA* and BLAST search analysis revealed the occurrence of a new *tcpA* allele, designated as *tcpA*^Env_Vm^. Phylogenetic analysis observed that this gene had sequence homology between 69.3 and 96.4 % when compared to other reported *tcpA* sequences, and could be categorized into a novel cluster clearly separated from other major *tcpA* clusters, including the classical, El Tor, Nandi, and Novais types. The novel *tcpA*^Env_Vm^ in *V. mimicus* strains showed the closest similarity with a couple of *V. cholerae* non-O1/non-O139 strains isolated from India and USA, and also with a *V. mimicus* strain (Acc. no. ACYV01000002) isolated from USA (Fig. 3). Most of the diversities observed among the *tcpA* alleles were in the carboxy-terminal half, but the amino-terminal region was almost conserved among the compared sequences. Comparative sequence analysis with the reference classical and El Tor strains of *V. cholerae* O1, showed that the *tcpA*^Env_Vm^ had 74% homology at the DNA level to that of the El Tor (N16961) and classical (O395) *tcpA*, with 40 and 43 substitutions, respectively, among 224 deduced amino acids of the *tcpA* gene (Fig. 3, Fig. S6). The Nandi and Novais types TcpA differed by 15 and 45 amino acid residues in comparison to that of the environmental *V. mimicus* of this study. Phylogenetic analysis also observed high sequence homology of one *V. mimicus* strain isolated from Brazil and another strain from China to the canonical TcpA of classical and El Tor O1 *V. cholerae*, respectively. However, these classical and El tor types TcpA of *V. mimicus* had 40 and 42 differences in amino acids, respectively, when compared to the TcpA^Env_Vm^.

**Fig. 3.**
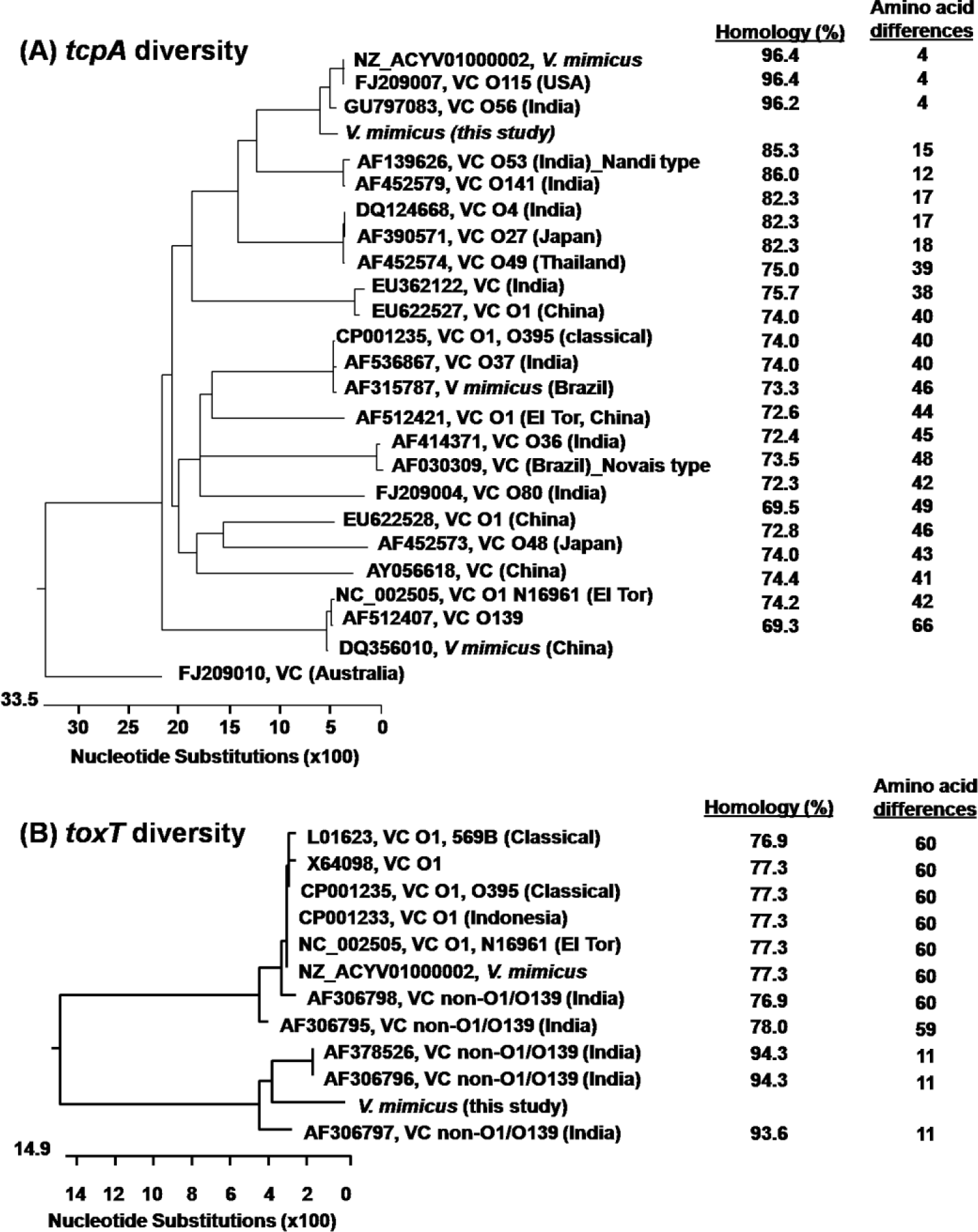
Genetic relatedness of *tcpA* and *toxT* genes among different serogroup strains of *V. cholerae* and *V. mimicus*. (A) The novel *tcpA* of *V. mimicus* of this study did not cluster in classical, El Tor, and other types of strains but formed a separate clade showing closeness to serogroups O56 and O115 strains of *V. cholerae*. (B) The novel *toxT* of *V. mimicus* of this study did not group into the major cluster comprising the *toxT* of *V. cholerae* O1 classical, El Tor, O139, non-O1/non-O139 and other *V. mimicus* strains but grouped into a separate cluster with atypical *toxT* reported in a few non-O1/non-O139 strains.

Similar to *tcpA* amplification, PCR using the conventional primers for *toxT* did not produce any amplicon for the environmental *V. mimicus* strains of this study. However, application of newly designed primers (Table S5), considering variations in classical, El Tor, and environmental types of *toxT*, successfully yielded specific amplicon of this gene in the study strains. Sequencing results showed identical sequence homology of *toxT* in all the environmental *V. mimicus* strains. Comparative analysis identified the presence of a new allele, with several unique substitutions, and 76.9-78.0 % homology among the deduced amino acid residues in comparison to the canonical *toxT* of the classical and El Tor *V. cholerae* strains (Fig. S7). Higher diversity was observed in the amino–terminal half of ToxT sequences when comparing those of the environmental *V. mimicus* and *V. cholerae* O1 strains. Phylogenetic analysis clearly differentiated *toxT* genes into two major clusters, one including the usual *toxT* commonly found in epidemic *V. cholerae* O1 strains and the other comprising the variant *toxT* identified in this study and several *V. cholerae* non-O1/non-O139 strains from India (Fig. 3). However, the variant ToxT in environmental *V. mimicus* strains was novel in terms of the acquired differences in 11 amino acid residues in comparison to that of the non-O1/non-O139 *V. cholerae* in the same phylogenetic cluster, and 59-60 amino acid residues with the canonical ToxT found in classical and El Tor *V. cholerae* O1.

### Competitive survival of *V. mimicus* in microcosm

In competition with the predominant estuarine vibrios, i.e., *V. cholerae* and *V. parahaemolyticus*, the inoculated *V. mimicus* strain could be cultured on TTGA agar up to 14, 45 and 55 days at 0.1, 3.5 and 11.5 ppt water salinities, respectively, in microcosm environment. The survival rate of *ctx*^+ve^ *V. mimicus* strain was comparable to a strain of epidemic *V. cholerae* O1. In contrast to a rapid decrease in culturable counts with time observed for *V. parahaemolyticus* at lower salinity (<5 ppt), the inoculated *V. mimicus* strain showed better potential to persist as culturable form at all the tested water salinities, representing their environmental habitats (Fig. S8).

### CT production capacity and virulence potential

All the environmental *V. mimicus* strains showed identical pattern for the major virulence related genes, including those of the predicted amino acid sequences in CTX prophages and TCP island, which indicated their probable functional capability to produce CT and virulence related proteins. Therefore, bead-ELISA was carried out following established conditions for both the El Tor and classical strains of *V. cholerae* O1 to check the functional CT production capacity and its variation, if any, among the environmental *V. mimicus* strains. Results showed that the CT production capacity varied among the environmental *V. mimicus* strains, and was better under the *in vitro* conditions favorable for the classical (LB, pH 6.6, 30 °C) strains than El Tor conditions (AKI, pH 7.4, 37 °C) for *V. cholerae* strains. Out of the six *V. mimicus* strains, one strain (Vm7) showed high CT production capacity (110 and 30 ng mL^−1^ in LB and AKI, respectively) while the toxin production was very low (0.1-0.5 ng mL^−1^) in others (Table 1).

SMA-based experiments produced results in congruence with CT production capacity for the *V. mimicus* strains (Table 1). Both live cells (10^6^ to 10^7^ CFU) and culture filtrates of the *V. mimicus* strain Vm7 producing high CT induced fluid accumulation and diarrhea in all the experimental mice, hence concluded to be enterotoxigenic. SMA score, representing fluid accumulation ratio, ranged between 0.083 and 0.090 (0.087 ± 003) for the high CT producing Vm7 strain. However, none of the experimental mice produced diarrhea when a low CT-producing *V. mimicus* strain Vm2 was administered at normal dose (10^7^ CFU) and even at higher dose (> 5×10^9^ CFU). The fluid accumulation ratio by this strain with attenuated CT production ranged between 0.065 and 0.070 (0.0068 ± 002), which was similar to that of the negative control (0.062 ± 002) (Table S9).

### Transcriptional analysis of genes associated with CT production

According to the results of bead-ELISA, the optimum culture condition for CT production, i.e., classical type condition using LB medium, was selected for transcriptional analysis of *ctxAB* and its known regulatory genes by qRT-PCR in the high (Vm7) and low (Vm2) CT producing environmental *V. mimicus* strains used in SMA *in vivo* experiments. As expected, a significantly lower transcription of *ctxA* in the low-CT producing strain in comparison to the high CT producer was observed. Similarly, significantly low-level transcription of *tcpA* and *toxT*, which are known to directly interact with CT production, was also observed. While checking the transcription of other genes in the ToxR regulon influencing CT production, the high and low transcription of *ctxA* was observed to be correlated with the mRNA transcription of *toxR, toxS*, and *tcpP* (Fig. 4). In the low CT-producing strain, the transcription of *ctxA, tcpA* and *toxT* genes was significantly lower by about 15- to 25-fold (P < 0.005), in comparison to those of the high CT producing strains. Similarly, *toxR* expression was also significant lower, about 4-fold (P <0.01), while both *toxS* and *tcpP* showed about 2- fold (P < 0.05) lower transcription. In the low CT producing strain, *tcpH* transcription was about 1.4-fold lower but not significant in comparison to the high CT producer. On the other hand, an opposite trend was observed for *hns* transcription; the high CT producing strain showed about 1.3-fold lower *hns* transcription, which was not significant, than the low CT producer.

**Fig. 4.**
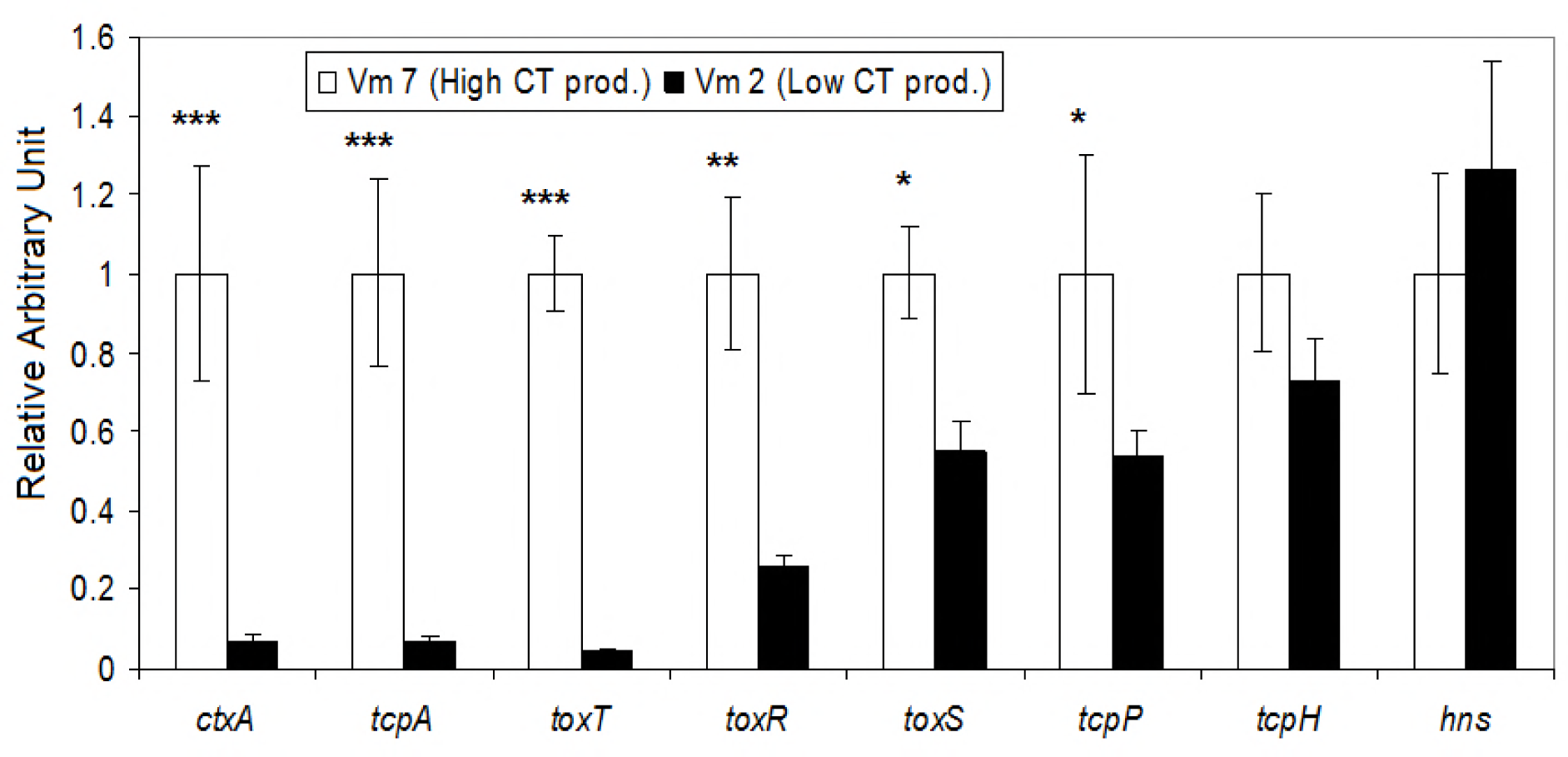
Variation in mRNA transcription of virulence and its regulated genes between a high and low CT-producing strains of *Vibrio mimicus*. Transcriptional levels of various virulence-related genes were analyzed by qRT-PCR. The relative transcriptional level of each gene was normalized with the housekeeping *recA* gene. The mRNA transcription level of each gene in a low CT-producing strain was compared with that of a high CT-producing strain. The transcriptional level of each virulence related gene of the high CT-producing strain was arbitrarily considered as 1 (Relative Arbitrary Unit). Statistically significant differences were calculated using the two-sample *t*-test. A P-value of <0.05 was considered as significant (*** = P <0.005; ** = P <0.01; * = P <0.05).

## DISCUSSION

Understanding the adaptive evolutionary mechanism of the CTXФ and *ctxAB* genes encoding the cholera toxin (CT) is highly important because of its direct relation to severe diarrhea such as cholera, which is causing health hazards throughout the world. In this aspect, through acquisition of toxigenic *ctxAB, V. mimicus* might play a salient role for its maintenance and propagation in the natural environment. Due to the lack of systematic surveillance of environmental and clinical samples, our knowledge of the occurrence and diversity of virulence genes associated with CT production in *V. mimicus* is very limited. In this study, the genetic traits and virulence potential of several estuarine strains of *ctx*^+ve^ *V. mimicus* were analyzed for better understanding of the role of this bacterium in the evolution of CTXФ, *ctxA*B, and related pathogenic factors.

### Novel *ctxAB* allele in CTX^ET^ prophage in environmental *V. mimicus* strains

Comparing the core and flanking regions of CTX prophage in *V. mimicus* with those of *V. cholerae* strains shows that some unique changes in amino acid residues, which were previously unidentified, with respect to the reference homologous genes have occurred in these *V. mimicus* strains. In the reference El Tor strain N16961, the phage integration site, characterized by the *attR* sequence followed by XerC and XerD, starts at 106 bp downstream of *ctxAB* intergenic region (27). Similar phage integration site starting at 106 bp downstream of *ctxAB* of CTXФ^ET^, which is different from pre-CTXФ^Env^ integration site, i.e., starting at 13 bp downstream of *zot*, has been observed in this study. Remarkably, comparison of amino acid residues with the known *ctxB* sequences has identified the presence of a novel *ctxB* variant (Table 2) in the *V. mimicus* strains. Phylogenetically, this newly discovered *ctxB* genotype 14 is distantly related to the El Tor genotype 3, but more closely related to the classical genotype 1 and Haitian genotype 7 (Fig. 2). However, sequencing results and comparison of *orfU* of CTX prophage in *V. mimicus* strains could identify its close homology with that of the El Tor type *V. cholerae* O1. Nonetheless, several unique changes in the amino acid residues within the first two (D1 and D2) of three domains of *orfU* (28) indicates the ongoing evolution of CTX prophage in environmental *V. mimicus* in parallel to those of the epidemic *V. cholerae* O1. According to Wang *et al*. (15), these polymorphic residues most likely interact with TolA and the ‘adsorption’ domains, associated with phage penetration.

Genome walking through hybridization and PCR demonstrated that the *ctx*^+ve^ *V. mimicus* strains actually contain one mature El Tor type prophage (CTXФ^ET^) with *ctxAB*, and two environmental type pre-CTX prophages (pre-CTXФ^Env^) without *ctxAB*. Existence of pre-CTXФ in some epidemic strains of *V. cholerae* O1 and O139 has been known (25). Integration of at least two types of CTXФ (El Tor and Environmental) within the genome of *V. mimicus* is an interesting novel observation. All of the *V. mimicus* strains in this study were observed to produce replicative forms of both CTXФ^ET^ and pre-CTXФ^Env^ when induced by mitomycinC in the culture filtrates, which was detected by PCRs after DNAse and RNAse treatment (data not shown). On the other hand, like the El Tor strains of *V. cholerae*, the environmental *V. mimicus* strains also harbored *rstC*, i.e., the RS1 element, which has been recently observed to promote diversity by the loss of CTX prophage and lysogenic immunity to make room for new CTX prophage to be integrated (29). Presence of *rstC* has been shown to increase *rstA* transcription and CTXФ production (17), which may influence *ctxAB* transcription and diversification. It is assumed that the El Tor strains possess greater ecological fitness than the classical strains. In comparison to the canonical El Tor type strains, the hybrid El Tor strains with classical type *ctxB* genotype is considered as more virulent than the El Tor CT producer (30). The acquisition of hybrid CTXФ^ET^ by the *V. mimicus* strains might have equipped them with greater evolutionary fitness, since these pathogenic strains can utilize the chance of becoming selectively enriched in the intestine of humans and animals.

### Diverse TCP and ToxT alleles in *V. mimicus* strains

Not only the CTX elements but also the TCP genes in *V. cholerae* can be mobilized by a generalized transduction (31). The sequence of the *tcpA* locus in the TCP element is known to be more divergent compared to other loci in the VPI (32). Similarly, by phylogenetic analysis we have observed a high diversity in *tcpA* sequences, compared to not only among the O1/O139 and non-O1/non-O139 strains of *V. cholerae* but also among the previously reported *ctx*^+ve^ *V. mimicus* strains. The observed homology of *tcpA* of *V. mimicus* and *V. cholerae*, showing as less as ca. 70% at nucleotide level, is in congruence with other studies analyzing strains belonging to different serotypes and biotypes of *V. cholerae* (33, 34). The environmental *ctx*^*+ve*^ *V. mimicus* strains contain a novel type of *tcpA* claimed as *tcpA*^Env_Vm^, which showed higher sequence homology to that of *V. cholerae* serogroups O56 and O115. Thus, *tcpA* of the environmental *V. mimicus* strains of this study might have acquired this gene from *V. cholerae* strains belonging to the O56 and O115 serogroups, and/or vice versa. This observation is not coherent with the previously reported *V. mimicus tcpA* genotypes, which were affiliated to the phylotgentic clades containing the canonical classical and El Tor strains of *V. cholerae* O1. Most likely, this diversity is a reflection of diversifying selection to CTXФ susceptibility during adaptation to the aquatic environment or host intestine. Sequencing results also indicate that *V. mimicus* strains contain a novel allele of *toxT*, affiliating with the atypical *toxT* of certain non-O1/non-O139 but not the canonical *toxT* of epidemic classical and El Tor O1 strains of *V. cholerae* (24).

Comparative analysis of *tcpA* sequences has clearly identified a high substitution rate in the carboxy-terminal half, encoding the exposed part of the TCP pilus on cell surface. Among the many differences between the present *tcpA*^Env_Vm^ allele and the *tcpA*^Cla^ allele (classical) is the c.187V>K substitution, which is shown to be correlated with increase in pilus-mediated autoagglutination in the context of *tcpA*^cla^ (35). In comparison, the amino-terminal region, encoding the basal part of the mature pilus structure, was observed to be more conserved among the *V. mimicus* and other strains compared. In case of *toxT*, the diversity in amino acid residuals in comparison to the reference strain was higher in the amino–terminal half, which is in accordance with a previous study (24). The relatively conserved carboxy-terminal half is known to determine the specificity of ToxT protein binding to DNA regulatory sites (24). Apart from acting as a virulence factor, TCP may also aid in the environmental persistence, e.g., biofilm formation on aquatic particles, and organisms, particularly, chitinous zooplankton (36). The occurrence of a new variant *tcpA*, with possible alterations in cell surface epitopes, among the toxigenic *V. mimicus* strains of the present study might be due to an adaptive evolutionary response to the changes in environmental niche.

### Variation in CT production, virulence potential, and its regulatory framework in *V. mimicus*

Results of bead-ELISA showed that CT production level in *V. mimicus* strains is preferentially induced under the *in vitro* growth conditions favorable for the classical *V. cholerae* O1 strains. Most of the *V. mimicus* strains did not cause fluid accumulation or diarrhea in experimental mice, in concordance with the very low CT production (<0.5 ng mL^−1^). Yet these environmental *V. mimicus* strains can be considered as potentially toxigenic because of their acquisition of genes related to pathogenic factors, including CT and TCP. This is reflected in at least one strain of this study producing considerable amount (>100 ng mL^−1^) of CT to induce fluid accumulation in suckling mice. We also cannot rule out the possibility that the current assay condition may not be suitable for inducing the CT production especially for low CT producing strains and that need to be further investigated using different culture conditions and growth media like M9-minimal medium. In case of *V. cholerae*, strains producing at least ∼20 ng mL^−1^ concentration of CT are known to cause fluid accumulation or diarrhea (37). Absence of any other potential virulence factors, e.g., TTSS, ST, ChxA and RTX indicates that the observed enterotoxicity in mice intestine is due to CT produced by the environmental *V. mimicus* strains. Interestingly, attenuation of some bacterial virulence factors has been attributed to the effect of repeated subculture *in vitro*, e.g., reduction of heat-labile enterotoxin (LT) in *E. coli*, and CT production in *V. cholerae*. (38; N. Chowdhury *et al*., unpublished). On the other hand, the higher CT production in one *V. mimicus* strain of this study may be due to its pre-exposure to the intestine of mammals, fish or any potential aquatic animal (39).

The studied *V. mimicus* strains showed resistance to polymixin B, ampicilin, cephalotin and reduced susceptibility to gentamicin. Resistance to polymixin B has been shown to be typical for the El Tor strains, while most of the O1 strains of both classical and El tor biotypes are usually resistant to amplicilin. When compared to the recent clinical strains from patients with cholera, the observed resistance to a few antimicrobials in the environmental *V. mimicus* strains is in congruence with the results obtained for *V. cholerae* O1 strains isolated from natural surface water (40). The widespread use of antimicrobials might have provided an additional selective pressure for the sporadic emergence of the multi-drug resistant *V. mimicus* strains.

Despite the presence of CTXФ and TCP element the variation in CT production is likely influenced by other genetic or physiological factors in *V. cholerae*. The expression of CT and TCP is activated by ToxT, which is regulated by the TcpP-TcpH-ToxR-ToxS complex of the ToxR regulon (22). In case of the high CT-producing *V. mimicus* strain, higher transcription of *ctxA* in conjunction with that of *tcpA* and *toxT* corroborates with the known ToxT-mediated genetic regulation influencing CT production in *V. cholerae*. Moreover, *ctxA* transcription was correlated with significant induction in the transcription of the upstream-regulatory genes *toxR/toxS*. In addition to ToxR regulon, the histone-like nucleoid structuring protein (H-NS) encoded by *hns*, a global prokaryotic gene regulator, has been shown to repress the transcription of several virulence genes including *toxT, ctxAB* and *tcpA* in *V. cholerae* (41). However, the variation in CT production in *V. mimicus* strains is not influenced by the H-NS since its transcription did not show considerable change in parallel to that of *ctxAB*. Therefore, the gene transcription results indicate that the ToxR/ToxS, in conjunction with ToxT, controls the CT production in *V. mimicus* strains. The ToxR regulon is thought to be controlled by environmental stimuli, such as temperature, pH and osmolarity (26). Hence, understanding the precise genetic and physiological mechanisms behind the very low or high level of CT production in environmental *V. mimicus* strains requires more extensive research, which is beyond the scope of this study.

### Probable role of *V. mimicus* in the evolution of CTXФ

The presence of the recombinase XerC and XerD binding sequences at both ends of the pre-CTXФ^Env^ and CTXФ^ET^ prophages support the phage-mediated integration events of these external genetic elements. The lack of CTXΦ element in some *tcpA*-positive O1 and non-O1/non-O139 strains supports the hypothesis that *tcpA* is acquired first and then integration of *ctxAB* genes happens during the evolution of pathogenic *V. cholerae* from their non-pathogenic progenitors (42). On the other hand, special forms of the CTXΦ family, designated as pre-CTXФ, do not carry *ctxAB* but contain other genes considered to be CTXΦ precursors (6). The step-wise occurrence of RS1-CTX^ET^-RS1-CTX^Env^-CTX^Env^ indicates an evolutionary signature of CTXФ insertion events in *V. mimicus*. The observed prevalence of novel types of *ctxAB, tcpA, toxT, and orfU*, indicates that CTX prophage on *V. mimicus* genome might have evolved independently of the 7^th^ pandemic El Tor clones, probably through independent integration of pre-CTXФ^Env^, in duplicate, and then a primeval CTXФ^ET^. The absence of RTX and TLC elements, which are usually located on the flanks of the CTX element in *V. cholerae* El Tor strains, also support this assumption for the environmental *ctx*^+ve^ *V. mimicus* strains. Similarly, these *V. mimicus* strains were devoid of the VSP I and VSP II genes cluster of pandemic El Tor strains of *V. cholerae*. Comparative whole genome sequence analysis also indicates horizontal transfer of virulence-related genes from an uncommon clone of *V. cholerae*, rather than the seventh pandemic strains, may have generated the pathogenic *V. mimicus* strain carrying *ctx* genes (9, 43). This is further supported by the observations of this study showing existence of five heptamer (TTTTGAT) repeats in the promoter region of *ctxAB* in *V. mimicus*, a characteristic genomic signature of the classical O1 strain isolated during 1960s, while the El Tor O1 strains, isolated since 1970s onward, contained four heptamer repeats (27). Therefore, the evolution of different types of CTXФ not only involves their integration into the epidemic strains of *V. cholerae* O1 but also environmental *tcp*^*+ve*^ *V. mimicus*. Based on the results of this study and previous reports of others, a hypothetical evolutionary map for the genomic drift associated with pathogenic traits in *V. mimicus* and *V. cholerae* has been depicted in Fig. 5.

**Fig. 5.**
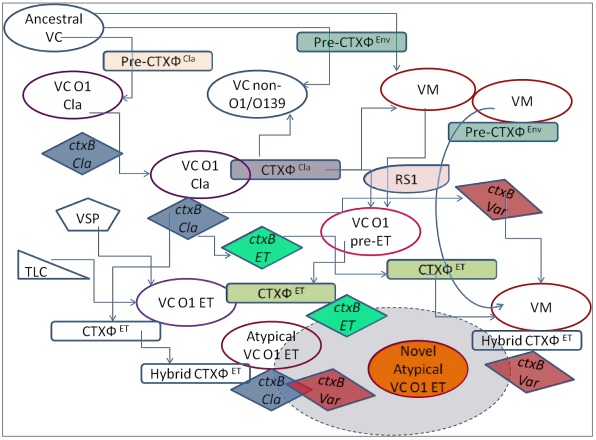
A hypothetical scenario of the evolution of CTX and variant virulence genes in *V. cholerae* and *V. mimicus*. Environmental *V. mimicus* may play a salient role by acting as an important reservoir of variant genes aiding the evolution. Line connectors with arrows indicate probable routes of origin of *V. mimicus* and *V. cholerae* strains containing *ctxB* variants. Bacteria, CTXФ, and *ctxB* gene are shown in different shapes, with solid line border. VM, VC, ET, Cla, VSP, and TLC indicate *V. mimicus, V. cholerae*, El Tor, Classical, *Vibrio* Seventh Pandemic island, and Toxin Linked Cryptic element, respectively. In the bottom, the light blue oval, with dotted border, indicates an interactive environmental pool facilitating generation of new clones of atypical *V. cholerae* O1 El Tor and *V. mimicus* strains possessing variant *ctxB* gene.

Isolation of clonally related *V. mimicus* strains with almost identical PFGE pulsotypes during different months of the year indicates their unique ancestral origin and high adaptation capacity in the estuarine environment. The microcosm results also support this notion as it shows prolong persistence of the *ctx*^+ve^ *V. mimicus* strains at least in culturable form, similar to *V. cholerae*, which are usually observed to co-occur in estuaries. This kind of *V. mimicus* strains, therefore, probably serves a cryptic but important natural reservoir of the CTXФ, TCP and related virulence genes. In the aquatic environment, lytic phage mediated transfer of virulence gene from classical *V. cholerae* to *V. mimicus* strain may have also happened through natural transformation in aquatic microhabitats including chitinous surface and biofilm (44). The probable influence of environmental *V. mimicus* on the ongoing population shift of typical El Tor strains to hybrid El Tor strains carrying classical and variant type *ctxB* cannot be overruled. However, clinical and environmental surveys until now have mainly focused on the detection of *ctx*^+ve^ *V. cholerae* strains. Thus, the cryptic existence of pre-CTXФ in both the *V. cholerae* and *V. mimicus* populations, which might provide significant evolutionary signatures regarding evolution of CTXФ and TCP element, has been so far neglected.

### Conclusion

It can be inferred that certain clonally related environmental *V. mimicus* strains can act as reservoir of variant *ctxB*, designated as genotype 14, which is phylogenetically more close to the currently predominant genotypes 1 and 7 associated with cholera outbreaks worldwide. This study provides molecular insight into the virulence potential of *ctx*^+ve^ *V. mimicus* strains, which could potentially serve as reservoirs of not only novel or variant type of *ctxAB*, but also *tcpA, toxT*, and *orfU* in the estuarine environment. The genomic content of tandemly arranged multiple pre-CTXФ^Env^ and a CTXФ^ET^ with novel classical type *ctxAB* probably act as salient raw materials for the natural recombination events driving the evolution of virulence genes related to CT production. Though CT production in some of this kind of environmental *V. mimicus* strains can be naturally attenuated, they may be potentially toxigenic in favourable conditions and can instigate cholera-like diarrhea. The variation in CT production capacity in *V. mimicus* is shown here to be controlled by the ToxR regulon, which is influenced by the physicochemical changes in the environment. The cryptic existence of the virulence genes related to CT production in *V. mimicus* genome points out an unnoticed event in the evolutionary pathway of CTXФ ecology and cholera epidemiology. Systematic environmental surveillance of non-epidemic strains, including *V. mimicus* and *V. cholerae*, and their detail molecular genetic analysis would allow us better understanding the evolution of new variant *ctx* element, and CTXФ, as well as the genes that regulate them.

## MATERIALS AND METHODS

### Bacterial strains and their antimicrobial susceptibility

Six *ctx*^+ve^ *V. mimicus* strains (Table 1) were obtained from the culture collection of Environmental Microbiology Laboratory of ICDDR,B. These strains were isolated during post-monsoon and early-winter months in 2000 from the Karnaphuli River estuary, Bangladesh. The *ctx*^+ve^ strains were screened from 1600 presumptive *V. mimicus* colonies, grown on thiosulfate citrate bile salts sucrose (TCBS) agar after enrichment of environmental samples in alkaline peptone water (pH 8.0) (APW). All strains were grown in Luria–Bertani (LB) broth, and their identity was verified according to standard protocol (45). Strains stored as glycerol stock at −80°C were grown in APW and subsequently on TCBS agar (Difco), Gelatin Agar (Difco), and LB at 37°C whenever needed. Several reference strains of *V. cholerae*, i.e., N16961 and O395, representing the El Tor and classical biotypes, respectively, VCE233 and AS522, non-O1/non-O139 strains containing environmental and Calcutta type CTX prophages, SG6, a Type III Secretion System (TTSS)-positive non-O1/non-O139, GP156, a *stn*-positive O1 El Tor and C9, a *chxA*-positive non-O1/non-O139, and *V. mimicus* ATCC 33653^T^ were used as controls. Each of the *ctx*^+ve^ *V. mimicus* strains were examined for resistance to some commonly used antibiotics (Table 1) by disc diffusion method according to the Clinical and Laboratory Standards Institute (http://www.clsi.org) using Mueller-Hinton agar (Difco Laboratories, MI, USA) and commercially available discs (Oxoid, Hampshire, England).

### Pulsed-field gel electrophoresis (PFGE)

PFGE was performed according to the Pulse Net USA protocol (http://www.cdc.gov/pulsenet/protocols.htm) with slight modifications. Briefly, freshly grown of *V. mimicus* strains were embedded into 1% Seakem Gold agarose followed by lysis of the cells with 0.5 mg mL^−1^ Proteinase K (P8044-5G, Sigma) and 1% Sarcosine (Sigma) at 54°C for 1 h. Agarose blocks containing genomic DNA were digested with *Not*I and *Sfi*I (30 and 40 U, respectively; Takara Bio Inc, Otsu, Japan) using appropriate buffer at 37°C for 3 h. DNA fragments were electrophoresed in 1% pulsed-field certified agarose gel (BioRad) using a CHEF MAPPER (Bio-Rad). Gels were stained for 30 min, de-stained twice for 15 min each and images were captured using a Gel-Doc 2000 (Bio-Rad). Lambda ladder (Bio-Rad) was used as a molecular mass standard. The PFGE fingerprints were analyzed by Fingerprinting II software (Bio-Rad).

### Colony blot hybridization of virulence related genes

DNA probes for colony blot hybridization included the major toxigenic factors, e.g., cholera toxin (*ctxA*), zonula occuldens toxin (*zot*, part of CTX phage), RS1 element (*rstC*) and *Vibrio* Seventh Pandemic island (VSP I and II, marker of present pandemic O1 El Tor biotype), TLC element, and other known virulence genes of *V. cholerae*, namely, *vcsN2,chxA, stn*, and *rtxA*, encoding Type III secretion system, cytotoxic cholix toxin, heat stable enterotoxin, and repeat in toxin, respectively. DNA templates of reference *V. cholerae* strains were subjected to PCR targeting the above mentioned virulence related genes. Standard PCR reaction mixture was prepared, applying primers for the toxigenic genes as mentioned in Table S5. PCR amplified genes were labeled by random priming with [α-^32^P]-dCTP (370 MBq mmol^−1^) using Multi-Prime DNA Labeling System (GE Healthcare, Buckinghamshire, UK). Environmental *V. mimicus* strains were grown on nitrocellulose membrane, overlaid on LB agar at 37°C for 4-6 h, and subjected to colony blot hybridization following the procedure described by Yamasaki *et al*. (46). Radioactivity in the hybridized membrane was detected using BAS FLA-3000 system (Fuji film, Tokyo, Japan).

### PCR based typing of virulence genes and CTX phage element

Template DNA was prepared by standard boiling method and stored at −30°C until use. The mismatch amplification mutation assay (MAMA)-PCR (47) was employed to detect *ctxB* genotype in *V. mimicus* strains to define their potential of classical or El Tor type CT production. The presence of the RS1 element was determined by the *rstC* gene-based PCR (17). Genotypes of *rstR*, namely, classical, El Tor, Calcutta and environmental, were also determined by PCR using a newly designed primer set for the environmental type, and previously established protocols for others (19). The genotypes of *tcpA*, belonging to the VPI, were also checked by PCR using previously established methods (48). Details of the primers and PCR conditions for screening these genes are mentioned in Table S5.

### Southern hybridization and PCR arrays to understand genetic organization of CTXФ

Southern hybridization of *Bgl*I and *Bgl*II digested gDNA of *V. mimicus* strains were carried out with probes for selected virulence related genes, including *ctxAB, rstR*^ET^, *rstR*^Env^, and *rstC*. Briefly, 5-μg aliquots of total gDNA were digested with the restriction enzymes using appropriate buffer and electrophoresed in 0.8% pulsed-field certified agarose gel (BioRad) using a CHEF MAPPER (BioRad). Once separated, the gDNA fragments were subjected to Southern transfer and blotted onto nylon membranes (Hybond-N^+^; Amersham). The genomic blots were hybridized with the gene probes, labeled by random priming with [α-32P]-dCTP (370 MBq mmol^−1^), and autoradiographed as described previously (12). In order to verify the genetic organization of CTXФ and associated elements, a series of PCR arrays were performed using the forward and reverse allelic primers, with multiple combinations, of genes *rstC, rstR^ET^, rstR^Env^,ctxAB* and *orfU* as shown in Fig. 1 using respective primers (Table S5).

### Nucleotide sequencing and phylogenetic analysis of virulence genes diversity

*V. mimicus* strains were subjected to nucleotide sequencing analysis for several target virulence genes of CTXФ, including *ctxAB, orfU, rstR*, and associated flanking regions comprising *zot*, intergenic regions (ig-1 and ig-2), and of TCP element, namely, *tcpA* and *toxT*. Briefly, PCRs using primers (Table S5) targeting these virulence genes and flanking regions were conducted following standard protocols. The amplified products were purified using QIAquick Purification Kit (QIAGEN GmbH, Hilden, Germany), then cycle sequencing was carried out using BigDye Terminator v3.1 Cycle Sequencing Kit according to the manufacturer’s instruction (Applied Biosystems). Afterwards, a further purification was done using CleanSEQ (Agencourt Bioscience), and nucleotide sequences were determined by an ABI PRISM 3100 Avant Genetic Analyzer (Applied Biosystems). The obtained gene sequences were assembled and aligned by DNA Lasergene software (DNASTAR, WI, USA). Homology search was performed using the BLAST program (http://blast.ncbi.nlm.nih.gov/Blast.cgi), and the nucleotide and deduced amino acid sequences were compared with published genes. Phylogenetic tree was constructed using ClustalW algorithm to understand the genetic lineage and sequence diversity within the study strains and other representative sequences of target gene of *V. mimicus* and *V. cholerae* published in GenBank.

### Microcosm experiments

Microcosm experiments were conducted to understand the competitive survival of *ctx*^+ve^ *V. mimicus* with predominant vibrios in water. Surface water samples, collected at the three isolation sites of *ctx*^+ve^ *V. mimicus* strains in the Karnaphuli estuary, with different salinities, i.e., 11.5, 3.5 and 0.1 ppt, and pH between approx. 7.6 and 8.0, were filter sterilized. Three microcosm sets, representing the isolation sites environment, were prepared in triplicate, each with 250 mL of sterile estuarine water in glass conical flasks (500 mL). One representative strain of each of *ctx*^+ve^ *V. mimicus, V. cholerae* O1, and *V. parahaemolyticus*, isolated from the same estuary, was added at ∼10^5^ CFU mL^−1^ to each microcosm and incubated at 25 °C. At regular intervals 100 μL sample was plated on Tauracholate Tellurite Gelatin Agar (TTGA, pH 7.5) and culturable vibrios were enumerated in triplicate following standard procedures. Colonies of the three species were differentiated according to their different size and morphology, biochemical test for sucrose utilization, and serology with specific antiserum for *V. cholerae* O1. Median (*n* = 3) counts of the culturable populations of each *Vibrio* species were compared.

### Measuring CT production by bead enzyme-linked immunosorbent assay (bead-ELISA)

The *ctx*-positive *V. mimicus* strains were grown in AKI-medium (pH 7.4) and Luria broth (L-broth, pH 6.6) (Difco, KS, USA) for 12 h at 37 and 30 °C, respectively, to compare their CT production in conditions favorable for the El Tor and classical strains of *V. cholerae* (31, 49). Subsequently, the OD_600_ nm of the bacterial cultures were adjusted to 1.0, followed by 100-fold dilution in respective media and incubation at stationary and shaking conditions, for 4 h each, at 180 rpm (49). The cell free supernatant (CFS) of each culture was prepared by centrifugation at 12,000 x*g* for 10 min followed by filtration through 0.22 μm filter (IWAKI, Tokyo, Japan). The CFS from each culture was diluted 10, 100 and 500 times with phosphate buffered saline (10 mM NaCl, pH 7.0) and the produced CT was measured by bead-ELISA. Purified CT was obtained following methodology described by Uesaka *et al.* (50) and used as controls for known concentration. Preparation of polyclonal rabbit antisera against CT, conjugation of Fab’ of antitoxin IgG with horseradish peroxidase, and estimation of CT secreted by each strain bead-ELISA were done according to Oku *et al*. (51). All experiments were done in triplicate.

### Detection of pathogenic potential *in vivo*

Two strains of *ctx*^+ve^ *V. mimicus* showing similar PFGE pulsotype but producing high and low CT, as detected by ELISA, were selected for evaluating enterotoxigenic potential *in vivo*. Suckling mice assay (SMA), using three-day-old Swiss albino suckling mice, was performed according to standard procedures (52). Briefly, an aliquot (0.1 mL) of freshly grown bacterial culture in LB medium, and also its filtrate (using 0.2 μm filter), was mixed with Evans Blue (0.01%, w/v) and intragastrically inoculated into each suckling mouse. Approximately 10^7^ CFU was inoculated as normal dose, however, higher and lower dose for low and high CT producer, respectively, were also administered. After 6 h of incubation, their intestines were removed, pooled and weighed. Fluid accumulation score in SMA was expressed as the ratio of weight of the intestine to the remaining body weight and a ratio of ≥ 0.08 was considered as positive. Culture filtrates of the reference strains of *ctx*^+ve^ *V. cholerae* O1 (O395), and *ctx*^*-ve*^ *V. mimicus* (ATCC 33653^T^) were used as positive and negative controls, respectively. Pathogenic potential of each strain was verified using five and three mice for live cells and culture filtrates, respectively.

### RNA isolation and qRT-PCR assay

The *ctx*^+ve^ *V. mimicus* strains, expressing high and low CT, were freshly grown up to the mid-logarithmic phase (∼10^8^ CFU mL^−1^) in LB medium following the classical condition of CT production (49). Total RNA was extracted and purified using Trizol reagent (Gibco-BRL, NY) according to the manufacturer’s instructions. The qRT-PCR assay was carried out with the primers and probes for genes, namely *ctxA, tcpA, toxT, toxR, toxS, tcpP, tcpH* and *hns*, which are known to regulate CT production and a housekeeping *recA* gene as an internal control (Table S5) following the TaqMan probe method. Each probe was labeled with FAM and TAMRA as 5′-reporter, and 3′-quencher dyes, respectively. Reverse transcription for cDNA synthesis from RNA template (1 μg) was carried out using the quick RNA-cDNA kit (Applied Biosystems Inc., CA) according to the manufacturer’s instruction. Real-time PCR was carried out using the amplified cDNA and TaqMan Gene Expression master mix containing each set of primer and probe (Applied Biosystems Inc.). PCR conditions were 50 °C for 2 min, 95 °C for 10 min and 40 cycles, each having 95 °C for 15 sec and 60 °C for 1 min, in an ABI PRISM 7000 sequence detection system (Applied Biosystems Inc.). The relative transcription in comparison with the internal control was analyzed according to Hagihara *et al*. (53).

### Statistical analysis

Statistica (ver. 10.0, StatSoft, Oklahoma, USA) was used to explore the differences between the mean values applying Student’s two-sample *t*-test. A *p*-value of <0.05 was considered as significant.

## ACKNOWLEDGEMENTS

This research was supported by the Osaka Prefecture University under the Monbukagakusho:MEXT scholarship and JASSO fellowship programs. We appreciate the technical support of the environmental surveillance team of icddr,b. The thoughtful suggestions received from Prodyot Kumar Basu Neogi, ex-scientist of icddr,b, are gratefully remembered. icddr,b is thankful to the Governments of Bangladesh, Canada, Sweden and the UK for providing core/unrestricted support.

## Author contributions

SBN and NC designed and performed laboratory experiments and participated in data analysis. SY and GBN coordinated the experiments and analyzed the data. ZHM and MSI performed field studies. SPA, MA and AH helped design the study and participated in laboratory experiments. SBN, NC and SY wrote the draft of manuscript. All authors read and approved the final manuscript.

## SUPPLEMENTARY INFORMATION

**Fig. S1.** PFGE analysis of environmental *ctx*^+ve^ and reference (ATCC) strains of *V. mimicus* (VM), and *ctx*^+ve^ non-O1/non-O139 (VCE 233), O1 El Tor (VC N16961) and O1 classical (VC O395) strains of *V. cholerae*. Left gel image, PFGE profiles of undigested gDNA showing similar size of the two chromosomes of *ctx*^+ve^ VM and ATCC VM strains. The middle and right gel images, PFGE patterns of *Not*I- and *Sfi*I-digested gDNA of *ctx*^+ve^ VM and the reference strains. The *ctx*^+ve^ VM strains were clonal but differing in 1-2 bands, indicated by arrows. *Not*I- and *Sfi*I-digested gDNA of *ctx*^+ve^ VM strains generated two (a and b) and three (I, II and III) PFGE profiles. Taken together, four PFGE profiles (Ia, Ib, IIa, IIb, and IIIb) could be distinguished among the six *ctx*^+ve^ VM strains. MW, molecular weight, representing the lambda ladder (Bio-Rad).

**Fig. S2.** PCR detection of the *rstC* (RS1), *rstR*^El^ ^Tor^, *rstR*^Calc^, *rstR*^Cla^ and *rstR*^Env^ genes, and the presence or absence of *ctxAB* in the El Tor type CTXФ and environmental type CTXФ in *V. mimicus* (Vm) strains. *V. cholerae* (Vc) strains belonging to O1 El Tor (N16961), O1classical (O395), and non-O1/O139 (AS522 and VCE233) were used as controls. Environmental *V. mimicus* strains were positive for *rstC* (RS1 element), *rstR*^El^ ^Tor^, and *rstR*^Env^ genes but did not contain *rstR*^Calc^ and *rstR*^Cla^ genes. Similar to a *V. cholerae* non-O1/O139 strain, VCE233, all of the environmental *V. mimicus* strains contained *ctxAB* in the El Tor type CTXФ but did not possess any *ctxAB* in the environmental type CTXФ.

**Fig. S3.** Probable genetic organization of El tor and environmental types of CTXФ, and RS1 element, deduced by comparison of the restriction map of the marker genes, i.e., *ctx, rstR^El^ Tor, rstR^Env^, and rstC*, respectively, in *V. mimicus* strains. Top panel: autoradiographed images of gDNA, of *V. mimicus* strains, digested by restriction enzymes (*Bgl*I or *Bgl*II) and detected by ^32^P-labelled PCR products of the marker genes. Bottom panel: a schematic diagram with location ofRS1, RS2 and Core of the CTX prophages, with lines (filled and dotted) showing the distance between the restriction sites, and bars mimicking the results of Southern hybridization using different probes. Taken together, the results indicated an array of RS1 - CTXФ^El^ ^Tor^ ^(ET)^ (with *ctxAB*) - RS1 - CTXФ^Env^ (without *ctxAB*) - CTXФ^Env^ (without *ctxAB*).

**Fig. S4.** Comparative variations in deduced amino acids of *orfU* gene sequences in selected *V. mimicus* and *V. cholerae* strains. Sequences were aligned by ClustalW algorithm. Amino acid positions are shown as a heading scale. Strain details are shown on the right border at each row. In comparison to *V. mimicus* strain in this study, only mismatched amino acids of *orfU* genes in other selected strains are shown while their identical amino acids are indicated by dots.

**Table S5.** Primers and probes used in this study.

**Fig. S6.** Genetic diversity of amino acid residues in the novel variant *tcpA* in *V. mimicus* strains of this study in comparison to that of the selected reference strains of *V. cholerae.* Sequences were aligned by ClustalW algorithm. Amino acid positions are shown as a heading scale. Strain details are shown on the right border at each row. In comparison to *V. mimicus* strain in this study, only mismatched amino acids of *tcpA* genes of other selected strains are shown while their identical amino acids are indicated by dots.

**Fig. S7.** Variation in amino acid residues in the novel variant *toxT* in *V. mimicus* strains of this study in comparison to other *toxT* genes in selected reference strains of *V. cholerae.* Sequences were aligned by ClustalW algorithm. Amino acid positions are shown as a heading scale. Strain details are shown on the right border at each row. In comparison to *V. mimicus* strain in this study, only mismatched amino acids of *toxT* genes in other selected strains are shown while their identical amino acids are indicated by dots.

**Fig. S8.** Competitive survival of *ctx*^+ve^ *V. mimicus* (Vm2), *V. cholerae* O1 and *V. paraheamolyticus* strains co-cultured in microcosm water having different salinities and pH. Filter sterilized water of three estuarine sites where *V. mimicus* strains were isolated was used in microcosm. Water samples of sites 1, 2 and 3 had salinity of 11.5, 3.5 and 0.1 ppt, respectively, and pH of 8.0, 7.7, and 7.6, respectively.

**Table S9:** Suckling mice assay showing enterotoxigenic potential of the *ctx*^+ve^ *V. mimicus* strains.

## REFERENCES

1) Safa A, Nair GB, Kong RY. 2010. Evolution of new variants of Vibrio cholerae O1. Trends Microbiol 18:46–54.

2) Rashed SM, Mannan SB, Johura FT, Islam MT, Sadique A, Watanabe H, Sack RB, Huq A, Colwell RR, Cravioto A, Alam M. 2012. Genetic characteristics of drug-resistant Vibrio cholerae O1 causing endemic cholera in Dhaka, 2006-2011. J Med Microbiol 61:1736–1745.

3) Saidi SM, Chowdhury N, Awasthi SP, Asakura M, Hinenoya A, Iijima Y, Yamasaki S. 2014. Prevalence of Vibrio cholerae O1 El Tor variant in a cholera-endemic zone of Kenya. J Med Microbiol 63:415–420.

4) Adewale AK, Pazhani GP, Abiodun IB, Afolabi O, Kolawole OD, Mukhopadhyay AK, Ramamurthy T. 2016. Unique clones of Vibrio cholerae O1 El Tor with Haitian Type ctxB allele implicated in the recent cholera epidemics from Nigeria, Africa. PLoS One 11:e0159794.

5) Faruque SM, Rahman MM, Asadulghani, Nasirul KMI, Mekalanos JJ. 1999. Lysogenic conversion of environmental Vibrio mimicus strains by CTXФ. Infect Immun 67:5723–5729.

6) Boyd EF, Moyer KE, Shi L, Waldor MK. 2000. Infectious CTXФ and the vibrio pathogenicity island prophage in Vibrio mimicus: evidence for recent horizontal transfer between V. mimicus and V. cholerae. Infect Immun 68:1507–1513.

7) Bi K, Miyoshi SI, Tomochika KI, Shinoda S. 2001. Detection of virulence associated genes in clinical strains of Vibrio mimicus. Microbiol Immunol 45:613–616.

8) Islam MS, Rahman MZ, Khan SI, Mahmud ZH, Ramamurthy T, Nair GB, Sack RB, Sack DA. 2005. Organization of the CTX prophage in environmental isolates of Vibrio mimicus. Microbiol Immunol 49:779–784.

9) Wang D, Wang H, Zhou Y, Zhang Q, Zhang F, Du P, Wang S, Chen C, Kan B. 2011. Genome sequencing reveals unique mutations in characteristic metabolic pathways and the transfer of virulence genes between Vibrio mimicus and Vibrio cholerae. PLoS One 6:e21299.

10) Dziejman M, Balon E, Boyd D, Fraser CM, Heidelberg JF, Mekalanos JJ. 2002. Comparative genomic analysis of Vibrio cholerae: genes that correlate with cholera endemic and pandemic disease. Proc Natl Acad Sci USA 99:1556–1561.

11) Chatterjee S, Ghosh K, Raychoudhuri A, Chowdhury G, Bhattacharya MK, Mukhopadhyay AK, Ramamurthy T, Bhattacharya SK, Klose KE, Nandy RK. 2009. Incidence, virulence factors, and clonality among clinical strains of non-O1, non-O139 Vibrio cholerae isolates from hospitalized diarrheal patients in Kolkata, India. J Clin Microbiol 47:1087–1095.

12) Awasthi SP, Asakura M, Chowdhury N, Neogi SB, Hinenoya A, Golbar HM, Yamate J, Arakawa E, Tada T, Ramamurthy T, Yamasaki S. 2013. Novel cholix toxin variants, ADP-ribosylating toxins in Vibrio cholerae non-O1/non-O139 strains, and their pathogenicity. Infect Immun 81:531–541.

13) Wang D, Wang X, Li B, Deng X, Tan H, Diao B, Chen J, Ke B, Zhong H, Zhou H, Ke C, Kan B. 2014. High prevalence and diversity of pre-CTXΦ alleles in the environmental Vibrio cholerae O1 and O139 strains in the Zhujiang River estuary. Environ Microbiol Rep 6:251–258.

14) Waldor MK, Rubin EJ, Pearson GD, Kimsey H, Mekalanos JJ. 1997. Regulation, replication, and integration functions of the Vibrio cholerae CTXФ are encoded by region RS2. Mol Microbiol 24:917–926.

15) Moyer KE, Kimsey HH, Waldor MK. 2001. Evidence for a rolling-circle mechanism of phage DNA synthesis from both replicative and integrated forms of CTXФ. Mol Microbiol 41:311–323.

16) Faruque SM, Asadulghani, Kamruzzaman M, Nandi RK, Ghosh AN, Nair GB, Mekalanos JJ, Sack DA. 2002. RS1 element of Vibrio cholerae can propagate horizontally as a filamentous phage exploiting the morphogenesis genes of CTXФ. Infect Immun 70:163–170.

17) Davis BM, Kimsey HH, Kane AV, Waldor MK. (2002) A satellite phage-encoded antirepressor induces repressor aggregation and cholera toxin gene transfer. EMBO J 21:4240–4249.

18) Basu A, Mukhopadhyay AK, Garg P, Chakraborty S, Ramamurthy T, Yamasaki S, Takeda Y, Nair GB. 2000. Diversity in the arrangement of the CTX prophages in classical strains of Vibrio cholerae O1. FEMS Microbiol Lett 182:35–40.

19) Bhattacharya T, Chatterjee S, Maiti D, Bhadra RK, Takeda Y, Nair GB, Nandy RK. 2006. Molecular analysis of the rstR and orfU genes of the CTX prophages integrated in the small chromosomes of environmental Vibrio cholerae non-O1, non-O139 strains. Environ Microbiol 8:526–534.

20) Faruque SM, Tam VC, Chowdhury N, Diraphat P, Dziejman M, Heidelberg JF, Clemens JD, Mekalanos JJ, Nair GB. 2007. Genomic analysis of the Mozambique strain of Vibrio cholerae O1 reveals the origin of El Tor strains carrying classical CTX prophage. Proc Natl Acad Sci USA 104:5151–5156.

21) Nguyen BM, Lee JH, Cuong NT, Choi SY, Hien NT, Anh DD, Lee HR, Ansaruzzaman M, Endtz HP, Chun J, Lopez AL, Czerkinsky C, Clemens JD, Kim DW. 2009. Cholera outbreaks caused by an altered Vibrio cholerae O1 El Tor biotype strain producing classical cholera toxin B in Vietnam in 2007 to 2008. J Clin Microbiol 47:1568–1571.

22) Faruque SM, Albert MJ, Mekalanos JJ. 1998. Epidemiology, genetics, and ecology of toxigenic Vibrio cholerae. Microbiol Mol Biol Rev 62:1301–1314.

23) Skorupski K, Taylor RK. 1997. Control of the ToxR virulence regulon in Vibrio cholerae by environmental stimuli. Mol Microbiol 25:1003–1009.

24) Mukhopadhyay AK, Chakraborty S, Takeda Y, Nair GB, Berg DE. 2001. Characterization of VPI pathogenicity island and CTXФ prophage in environmental strains of Vibrio cholerae. J Bacteriol 183:4737–4746.

25) Mantri CK, Mohapatra SS, Colwell RR, Singh DV. 2010. Sequence analysis of Vibrio cholerae orfU and zot from pre-CTXΦ and CTXΦ reveals multiple origin of pre-CTXΦ and CTXΦ. Environ Microbiol Rep 2:67–75.

26) Tenover FC, Arbeit RD, Goering RV, Mickelsen PA, Murray BE, Persing DH, Swaminathan B. 1995. Interpreting chromosomal DNA restriction patterns produced by pulsed-field gel electrophoresis: criteria for bacterial strain typing. J Clin Microbiol 33:2233–2239.

27) Halder K, Das B, Nair GB, Bhadra RK. 2010. Molecular evidence favouring step-wise evolution of Mozambique Vibrio cholerae O1 El Tor hybrid strain. Microbiology 156:99–107.

28) Heilpern AJ, Waldor MK. 2003. pIIICTX, a predicted CTXФ minor coat protein, can expand the host range of coliphage fd to include Vibrio cholerae. J Bacteriol 185:1037–1044.

29) Kamruzzaman M, Robins WP, Bari SM, Nahar S, Mekalanos JJ, Faruque SM. 2014. RS1 satellite phage promotes diversity of toxigenic Vibrio cholerae by driving CTX prophage loss and elimination of lysogenic immunity. Infect Immun 82:3636–3643.

30) Naha A, Chowdhury G, Ghosh-Banerjee J, Senoh M, Takahashi T, Ley B, Thriemer K, Deen J, Seidlein LV, Ali SM, Khatib A, Ramamurthy T, Nandy RK, Nair GB, Takeda Y, Mukhopadhyay AK. 2013. Molecular characterization of high-level-cholera-toxin-producing El Tor variant Vibrio cholerae strains in the Zanzibar Archipelago of Tanzania. J Clin Microbiol 51:1040–1045.

31) O’Shea YA, Boyd EF. 2002. Mobilization of the Vibrio pathogenicity island between Vibrio cholerae isolates mediated by CP-T1 generalized transduction. FEMS Microbiol Lett 214:153–157.

32) Sarkar A, Nandy RK, Nair GB, Ghose AC. 2002. Vibrio pathogenicity island and cholera toxin genetic element-associated virulence genes and their expression in non-O1 non-O139 strains of Vibrio cholerae. Infect Immun 70:4735–4742.

33) Kumar P, Thulaseedharan A, Chowdhury G, Ramamurthy T, Thomas S. 2011. Characterization of novel alleles of toxin co-regulated pilus A gene (tcpA) from environmental isolates of Vibrio cholerae. Curr Microbiol 62:758–763.

34) Li F, Du P, Li B, Ke C, Chen A, Chen J, Zhou H, Li J, Morris JG (Jr), Kan B, Wang D. 2014. Distribution of virulence-associated genes and genetic relationships in non-O1/O139 Vibrio cholerae aquatic isolates from China. Appl Environ Microbiol 80:4987–4992.

35) Kirn TJ, Lafferty MJ, Sandoe CM, Taylor RK. 2000. Delineation of pilin domains required for bacterial association into microcolonies and intestinal colonization by Vibrio cholerae. Mol Microbiol 35:896–910.

36) Reguera G, Kolter R. 2005. Virulence and the environment: a novel role for Vibrio cholerae toxin-coregulated pili in biofilm formation on chitin. J Bacteriol 187:3551–3555.

37) Ghosh-Banerjee J, Senoh M, Takahashi T, Hamabata T, Barman S, Koley H, Mukhopadhyay AK, Ramamurthy T, Chatterjee S, Asakura M, Yamasaki S, Nair GB, Takeda Y. 2010. Cholera toxin production by the El Tor variant of Vibrio cholerae O1 compared to prototype El Tor and classical biotypes. J Clin Microbiol 48:4283–4286.

38) Ram S, Khurana S, Singh RP, Khurana SB. 1992. Loss of some virulence factors of enterotoxigenic Escherichia coli on repeated subcultures. Indian J Med Res 95:284–287.

39) Tikoo A, Singh DV, Sanyal SC. 1994. Influence of animal passage on haemolysin and enterotoxin production in Vibrio cholerae O1 biotype El Tor strains. J Med Microbiol 40: 246–251.

40) Chakraborty S, Garg P, Ramamurthy T, Thungapathra M, Gautam JK, Kumar C, Maiti S, Yamasaki S, Shimada T, Takeda Y, Ghosh A, Nair GB. 2001. Comparison of antibiogram, virulence genes, ribotypes and DNA fingerprints of Vibrio cholerae of matching serogroups isolated from hospitalised diarrhoea cases and from the environment during 1997-1998 in Calcutta, India. J Med Microbiol 50:879–888.

41) Nye MB, Pfau JD, Skorupski K, Taylor RK. 2000. Vibrio cholerae H-NS silences virulence gene expression at multiple steps in the ToxR regulatory cascade. J Bacteriol 182:4295–4303.

42) O’Shea YA, Reen FJ, Quirke AM, Boyd EF. 2004. Evolutionary genetic analysis of the emergence of epidemic Vibrio cholerae isolates on the basis of comparative nucleotide sequence analysis and multilocus virulence gene profiles. J Clin Microbiol 42:4657–4671.

43) Hasan NA, Grim CJ, Haley BJ, Chun J, Alam M, Taviani E, Hoq M, Munk AC, Saunders E, Brettin TS, Bruce DC, Challacombe JF, Detter JC, Han CS, Xie G, Nair GB, Huq A, Colwell RR. 2010. Comparative genomics of clinical and environmental Vibrio mimicus. Proc Natl Acad Sci USA 107:21134–21139.

44) Udden SM, Zahid MS, Biswas K, Ahmad QS, Cravioto A, Nair GB, Mekalanos JJ, Faruque SM. 2008. Acquisition of classical CTX prophage from Vibrio cholerae O141 by El Tor strains aided by lytic phages and chitin-induced competence. Proc Natl Acad Sci USA 105:11951–11956.

45) Davis BR, Fanning GR, Madden JM, Steigerwalt AG, Bradford HB, Smith HL, Brenner DJ. 1981. Characterization of biochemically atypical Vibrio cholerae strains and designation of a new pathogenic species, Vibrio mimicus. J Clin Microbiol 14:631–639.

46) Yamasaki S, Garg S, Nair GB, Takeda Y. 1999. Distribution of Vibrio cholerae O1 antigen biosynthesis genes among O139 and other non-O1 serogroups of Vibrio cholerae. FEMS Microbiol Lett 179:115–121.

47) Morita M, Ohnishi M, Arakawa E, Bhuiyan NA, Nusrin S, Alam M, Siddique AK, Qadri F, Izumiya H, Nair GB, Watanabe H. 2008. Development and validation of a mismatch amplification mutation PCR assay to monitor the dissemination of an emerging variant of Vibrio cholerae O1 biotype El Tor. Microbiol Immunol 52:314–317.

48) Rivera IN, Chun J, Huq A, Sack RB, Colwell RR. 2001. Genotypes associated with virulence in environmental isolates of Vibrio cholerae. Appl Environ Microbiol 67:2421–2429.

49) Iwanaga M, Yamamoto K, Higa N, Ichinose Y, Nakasone N, Tanabe M. 1986. Culture conditions for stimulating cholera toxin production by Vibrio cholerae O1 El Tor. Microbiol Immunol 30:1075–1083.

50) Uesaka Y, Otsuka Y, Lin Z, Yamasaki S, Yamaoka J, Kurazano H, Takeda Y. 1994. Simple method of purification of Escherichia coli heat-labile enterotoxin and cholera toxin using immobilized galactose. Microb Pathogenesis 16:71–76.

51) Oku Y, Uesaka Y, Hirayama T, Takeda Y. 1988. Development of a highly sensitive bead-ELISA to detect bacterial protein toxins. Microbiol Immunol 32:807–816.

52) Dean AG, Ching TC, Williams RG, Harden LB. 1972. Test for Escherichia coli enterotoxin in infant mice: application in a study of diarrhea in children in Honolulu. J Infect Dis 125:407–411.

53) Hagihara K, Nishikawa T, Isobe T, Song J, Sugamata Y, Yoshizaki K. 2004. IL-6 plays a critical role in the synergistic induction of human serum amyloid (SAA) gene when stimulated with proinammatory cytokines as analyzed with an SAA isoform real-time quantitative RT-PCR assay system. Biochem Bioph Res Co 314:363–369.

54) Marin MA, Vicente AC. 2012. Variants of Vibrio cholerae O1 El Tor from Zambia showed new genotypes of ctxB. Epidemiol Infect 140:1386–1387.

55) Zhang P, Zhou H, Kan B, Wang D. 2013. Novel ctxB variants of Vibrio cholerae O1 isolates, China. Infect Genet Evol 20:48–53.

